# Transposable element methylation state predicts age and disease

**DOI:** 10.1101/2024.03.15.585206

**Authors:** Francesco Morandini, Jinlong Y. Lu, Cheyenne Rechsteiner, Aladdin H. Shadyab, Ramon Casanova, Beverly M. Snively, Andrei Seluanov, Vera Gorbunova

## Abstract

Transposable elements (TEs) are DNA sequences that expand selfishly in the genome, possibly causing severe cellular damage. While normally silenced, TEs have been shown to activate during aging. DNA methylation is one of the main mechanisms by which TEs are silenced and has been used to train highly accurate age predictors. Yet, one common criticism of such predictors is that they lack interpretability. In this study, we investigate the changes in TE methylation that occur during human aging. We find that evolutionarily young LINE1s (L1s), the only known TEs capable of autonomous transposition in humans, undergo the fastest loss of methylation, suggesting an active mechanism of de-repression. We then show that accurate age predictors can be trained on both methylation of individual TE copies and average methylation of TE families genome wide. Lastly, we show that while old L1s gradually lose methylation during the entire lifespan, demethylation of young L1s only happens late in life and is associated with cancer.

## Introduction

Repetitive elements (REs) are DNA sequences found in high copy number in the genome^1^. Transposable elements (TEs), or selfish REs, are REs that have the ability to copy themselves and move to new genomic locations, either directly as DNA (DNA transposons) or through an RNA intermediate that is reverse-transcribed (LINEs, SINEs, LTRs). The selfish replication of TEs has led them to occupy a large portion of genomes (around 40% in mammals). TE activity is potentially highly detrimental to the individual, as random integrations can disable genes and even unsuccessful integration attempts can generate double stranded breaks^2^. Even further, TEs can produce cDNA copies that stimulate cytosolic DNA sensing pathways leading to inflammation^3–6^. Finally, TEs can disrupt normal gene regulatory networks by influencing the expression of nearby genes through their regulatory sequences^7^. Due to their pathogenic potential, TEs are kept under tight control by the host with multiple regulatory layers^8^. DNA methylation is one of the main ways by which cells silence TEs^9^. DNA methylation patterns are established in bulk during development and are then largely maintained throughout lifespan, although de-novo methylation and active demethylation still occur^10^. Prior studies in multiple organisms and tissues found that methylation patterns undergo a slow drift during aging, with many normally hypermethylated regions becoming less repressed^11–13^. At the same time, TEs have been shown to activate during aging in invertebrates, mice, human senescent cells, and certain cancers^2,14,15^. It thus seems possible that age-related alterations of DNA methylation could play a role in TE activation.

Aging clocks are statistical models trained to predict age and age-related phenotypes, including time to death^16^. In addition to predicting the age of samples of unknown age, for example in forensics, aging clocks have been used to study health conditions, lifestyles, genetic or pharmacological treatments that alter an organism’s biological age. Typically, age predictions are based on omic data types including gene expression^17,18^, protein abundance^19^, chromatin accessibility^20^ and most commonly, DNA methylation^21–26^. One common criticism of aging clocks deals with the difficulty in interpreting the biological meaning of observed changes in DNA methylation patterns. One strategy previously used to improve clock interpretability is to group clock CpGs into different modules corresponding to different biological processes^27,28^.

In this study, we explore the use of TE methylation as a biomarker of age and disease. First, we reanalyzed public human blood methylation data to determine the trajectory of TE methylation during aging, comparing evolutionarily young and old TEs. We then constructed age predictors for mice and humans. Lastly, we investigated associations between accelerated age prediction, and more generally loss of methylation at TEs, and disease.

## Results

### Data description

To investigate changes in RE methylation that occur during aging we collected publicly available human blood methylation array data. Later, we additionally investigate association between TE methylation and disease using the Women’s Health Initiative (WHI) BA23 dataset. The dataset characteristics are summarized in **Figure 1a**. All datasets were generated with the Illumina Infinium 450k array, which measures methylation at 485578 CpGs. We annotated array CpGs based on the type of RE and genic region (Exon, intron, promoter, 5’ UTR, 3’ UTR, intergenic) they lied within. Array CpGs were generally biased to genic regions, whereas complex repeats generally lie in intergenic regions or introns (**Supplementary figure 1a**). Nonetheless, 69426 CpGs were contained within REs, mainly LINEs, SINEs, LTRs, DNA transposons, and simple repeats, **Supplementary figure 1b**). While most RE CpGs were primarily intergenic and intronic (**Supplementary figure 1b**), simple repeats and low complexity regions were predominantly found in promoters.

**Figure 1:**
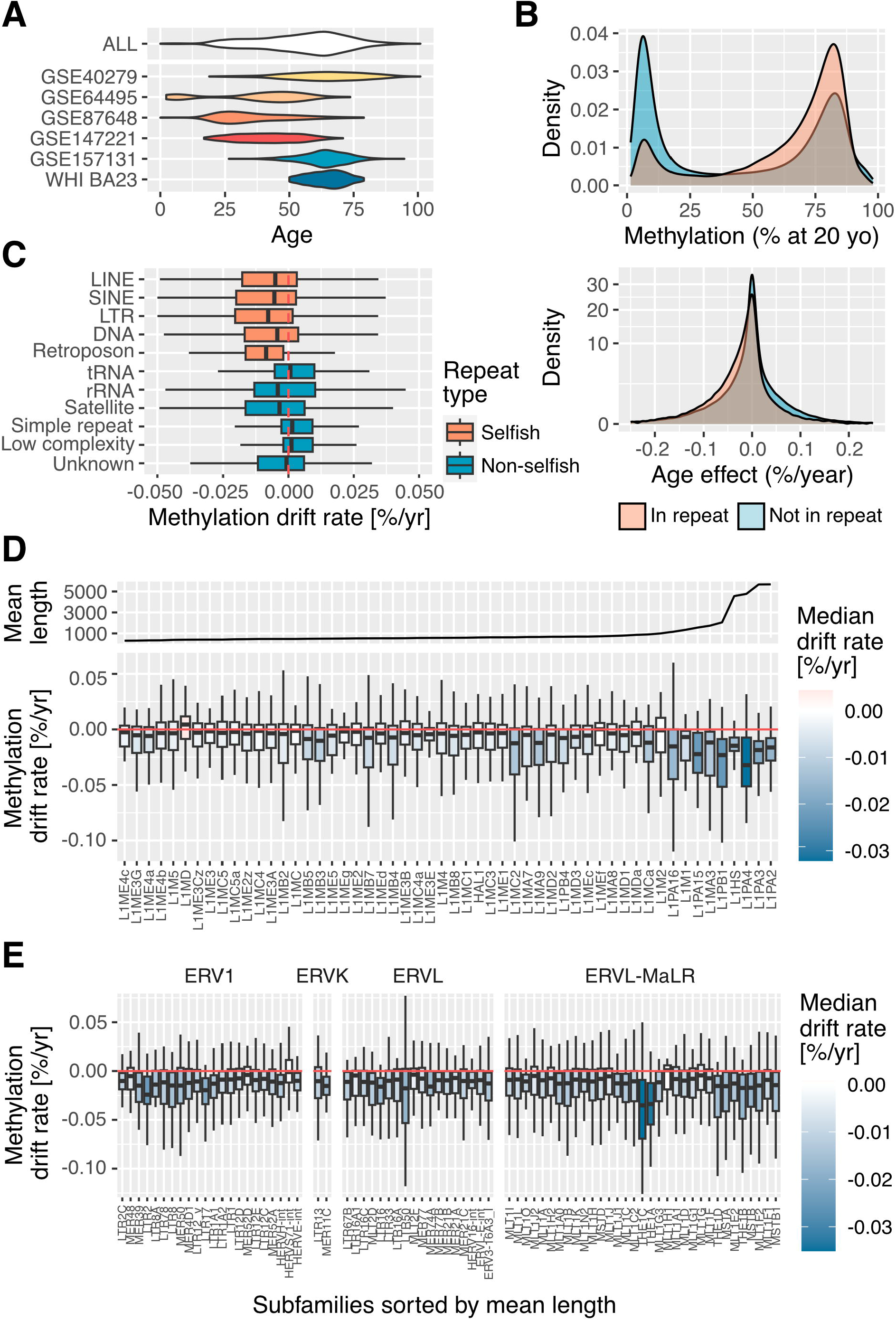
Transposons and particularly young L1s are biased towards losing methylation during aging. (A) Public human blood DNA methylation datasets and age distributions. (B) Youthful methylation level and age-related drift of CpGs in and outside of repetitive elements. (C) Methylation drift rate of CpG, grouped by major repeat class. Selfish (transposons) and non-selfish repeated grouped separately. (D) Methylation drift rate of CpG in L1s, grouped by family and sorted by average sequence length: a proxy of evolutionary age. Only families represented by 40 or more CpGs in the infinium array were shown (E) Methylation drift rate of CpG in LTRs, grouped by family and sorted by average sequence length. Only families represented by 40 or more CpGs in the infinium array were shown.

### Transposable elements and especially young L1s become derepressed during aging

Next, we investigated the age dynamics of RE and non-RE CpGs. We used limma^29^ to fit linear regression models to the methylation levels of all array CpGs including age, sex and the study of origin as independent variables (**Data file 1**). Patients with reported health conditions in the original studies were not included in the analysis, to initially focus on RE methylation changes that are associated with aging rather than disease. RE CpGs were hypermethylated in young individuals (20 years old), but were more likely to have decreased methylation in older individuals, compared to non-RE CpGs (**Figure 1b**). When investigating different classes of REs individually, we found that TEs (LINEs, SINEs, LTRs, DNA transposons, retroposons) were much more prone to losing methylation than non-selfish REs (tRNA, rRNA, satellites, simple repeats, low complexity regions. **Figure 1c**). We initially focused on L1s, since they are the only TEs known to be active and autonomous in humans^30^. Therefore, de-repression of L1s could be sufficient to cause cellular damage. Fortunately, most L1 copies are truncated, or have mutated over evolutionary time scales and are thus inactive^31,32^. Conversely, competent, evolutionarily young L1 copies are closer to 6000 bp long. We found an association between the average length of L1 families and their propensity to become demethylated with age (**Figure 1d**). The most extreme methylation loss was observed in L1HS, L1PA2, L1PA3 and L1PA4, which are the 4 youngest L1 families present in the human genome^31^. Older families were also generally prone to methylation loss, but to a much smaller extent. We then investigated other TE classes: among LTRs, families THE1A and THE1C showed the fastest methylation loss (**Figure 1e**). While not retrotransposition-competent, derepression of these families was shown to drive expression of oncogenes^7^. Most SINE and DNA transposons were also biased towards losing methylation during age, but the median drift rate was relatively small, and no particular family stood out. (**Supplementary figure 1c, d**)

### Demethylation of young L1s outpaces passive methylation loss

The difference in demethylation rate between young and old L1 could indicate that they become de-repressed by different means: de-repression of old L1s may be a result of global age-related methylation loss, which has been previously documented and is often attributed to imperfect maintenance of methylation marks by DNMT1^12^. Conversely, young L1s may actively de-repress by recruiting activating transcription factors at their 5’ UTR^33^. Alternatively, this discrepancy may be explained by differences between the CpG landscape of young and old L1 families. For example, young L1s have a higher CpG density, which is gradually lost over evolutionary time scales due to C-to-T mutations^34^, and CpG density has been shown to affect the rate of passive methylation loss^35,36^. Additionally, the initial (post-development) level of CpG methylation may affect the methylation drift rate simply because highly polarized states (e.g. fully methylated) can only lose methylation, while intermediate methylation states are able to both gain and lose methylation during aging. Thus, we modelled the average methylation drift rate of CpGs based on local CpG density, youthful methylation level and the interaction of the two (**Figure 2a**). This model explained 24.7% of age coefficient variation and confirmed prior reports that low CpG density associates with age-related methylation loss. Hypomethylated CpGs (< 20% methylated) were more likely to gain methylation during aging, but hypermethylated CpGs (> 80% methylated) were not particularly biased towards methylation loss. We then adjusted the previously calculated age coefficients with this information (**Data file 1**). These adjusted age coefficients should be interpreted as “the age drift rate of a given CpG, compared to what would be expected from the average CpG with the same local CpG density and youthful methylation level”. The adjusted coefficients of most TE families of all 4 major classes were close to zero or even slightly positive, meaning that their aging trajectory could be explained by the local CpG context and youthful methylation state and is likely a passive phenomenon (**Figure 2b, c, d, e**). Conversely, L1HS, L1PA2, L1PA3 and L1PA4 retained a higher-than-expected rate of methylation loss, reinforcing the hypothesis that their derepression may be, at least in part, an active process.

**Figure 2:**
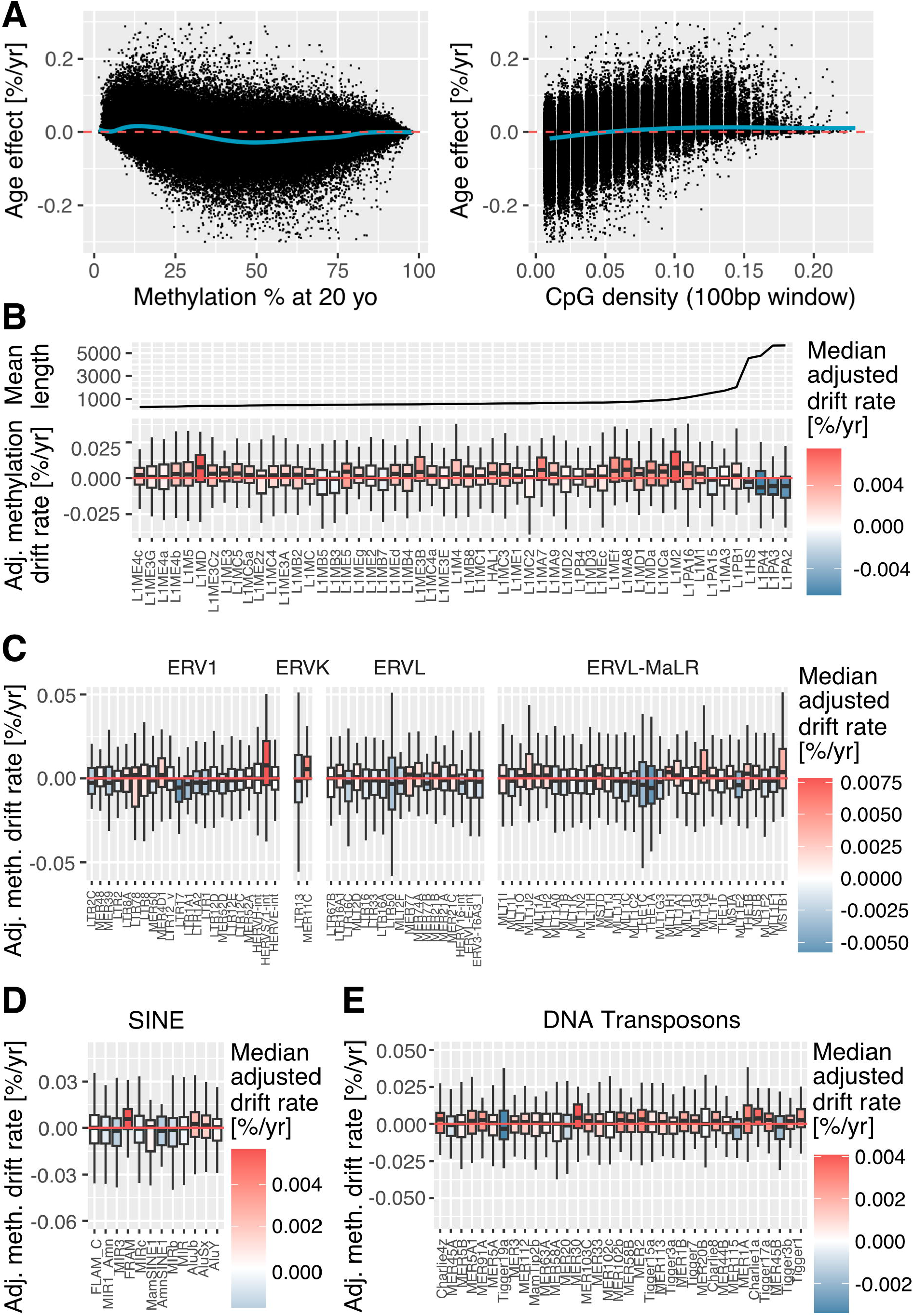
Age drift of TE CpGs compared to what is expected based on CpG density and youthful methylation level. (A) Trends of methylation drift based on youthful methylation levels. (B) Trends of methylation drift based on local CpG density. (C,D,E,F) Age coefficient of methylation at LINEs, LTRs, SINEs and DNA transposon CpGs after adjustment for CpG density and youthful methylation level. Only families represented by 40 or more CpGs in the infinium array were shown.

### TE methylation as an accurate and interpretable biomarker of age

Next, we investigated if the methylation state of TEs could be used to predict chronological age. Thus, we selected CpGs found in TEs (LINE, SINE, LTR, DNA transposons, ncpg=56352, **figure 3a**) and trained an elastic net model on a portion of our data (n=999), leaving out a portion of each dataset (n=248) and the entirety of GSE64495 (n=104) as external validation (**Figure 3b, c**). The coefficients are available in **data file 2**. This individual CpG TE clock was in both cases highly accurate (RMSE = 5.58, MAE = 2.96, r = 0.95 on GSE64495). We compared this performance with other state-of-the-art chronological age clocks and found that the individual CpG TE clock performed better than the Hannum and Horvath pan-tissue clocks but worse than Horvath Skin & Blood. Thus, the methylation state of individual CpGs within TEs can be used to construct a remarkably accurate clock.

**Figure 3:**
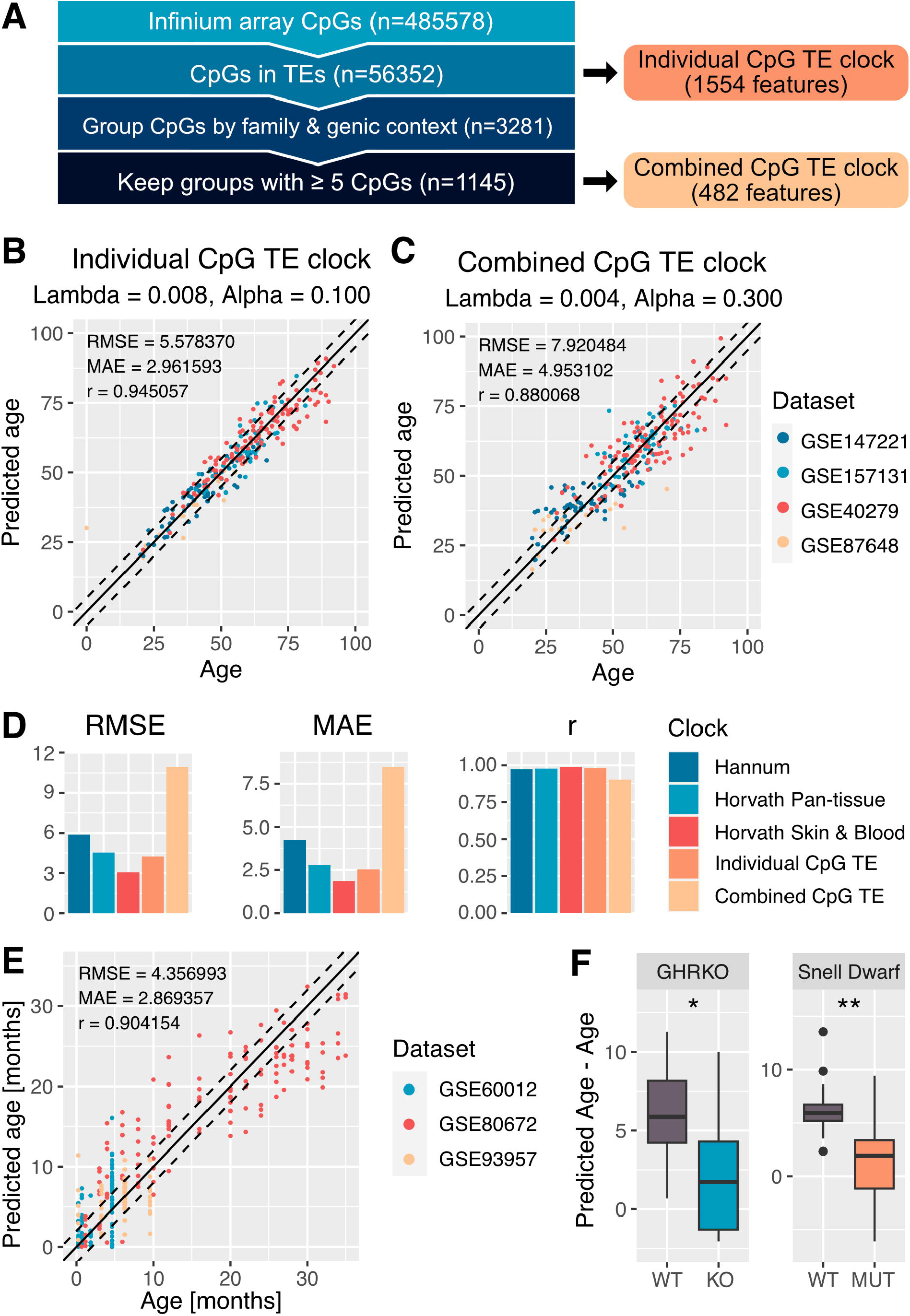
Construction of age biomarkers based on methylation of individual CpGs within TEs and genome-wide TE family methylation. (A) Feature construction strategy. (B) Test set performance of single CpG clock. (C) Test set performance of combined CpG clock. (D) Benchmark of individual and combined CpG clock against state-of-the art methylation clocks. The benchmark was performed on GSE64495, which was not included in the training set of any of the clocks shown. (E) Performance of a combined CpG clock trained on multi-tissue mouse RRBS data. (F) Age prediction on long lived mouse strains compared to matching controls.

While constructing a biomarker on a particular biological process such as TE derepression can indeed help with interpretability, further considerations should be made. Most importantly, transposons are disseminated everywhere in the genome, including near genes and very commonly in introns. Thus, while the state of methylation of a single TE CpG may be representative of the status of that TE copy, it may also be affected by the local chromatin context (for example, whether a nearby gene is transcribed or not). To further improve interpretability, we trained a new clock, this time on the average genome-wide methylation state of TE families, separating genic and intergenic TE copies. We chose not to completely discard genic TE copies because a sizeable portion of TEs, including some active L1s, is found in introns. Additionally, we only kept groups of at least 5 CpGs, to reduce the impact of the local regulatory context at each CpG and ensure that each feature could be interpreted as the global methylation of a given TE family (**Figure 3a**). Validation was again performed on a portion of each dataset (n=248) and the entirety of GSE64495 (n=104). The coefficients are available in **data file 2**. We were surprised to see that while performance of this Combined CpG TE clock was worse than that of the individual CpG TE clock, it was still satisfactory (**Figure 3b, c**). In particular, it still had an r of 0.90 when validated on the external dataset GSE64495.

Lastly, we applied the same combined CpG training strategy on reduced representation bisulfite sequencing (RRBS) data of multiple mouse tissues. Due to the limited data availability, the predictor was trained and validated using nested cross validation, once again only including wild-type, untreated mice (n=276). The coefficients are available in **data file 2**. This again yielded an accurate predictor, with r = 0.90 (**Figure 3e**). Thus, our feature construction strategy is successful on multiple sequencing platforms, tissues and organisms. We note that while the strategy is indeed successful across different species, generating a single TE-based biomarker for multiple species would be difficult, as TEs evolve very rapidly. For example, mice and humans have a very different number and set of active TEs^31,37^.

### Accelerated TE methylation age is associated with health status

Next, we investigated associations between age acceleration (the difference between predicted and chronological age) and health status. We tested our biomarkers on methylation data from the Women’s Health Initiative (WHI), a long-term study, deeply phenotyped among postmenopausal women. Specifically, we used data from substudy BA23, comprising 2175 women aged 50-79 years at baseline, of which ∼1070 developed coronary heart disease (CHD) during follow-up. We examined associations between age acceleration and time to death, diagnosis of any cancer, and CHD using Cox regression, including chronological age as a covariate. Accelerated aging according to the individual CpG TE clock was significantly associated with higher risk for all three outcomes (**Figure 4a**). Age acceleration according to PhenoAge^24^, an aging biomarker trained on clinical phenotypes rather than chronological age alone, had similar associations with risk of cancer and mortality as our individual CpG TE clock. Increased CHD risk, however, was most associated with age acceleration according to PhenoAge. Our combined CpG TE clock, on the other hand showed no significant associations with cancer or CHD risk, but was still associated with risk of death. We suspect this may be due to the decreased accuracy of this predictor, which relies on genome-wide methylation features. We additionally tested our mouse RRBS clock on data from Petkovich et al. comprising long-lived growth hormone KO (GHRKO) and Snell dwarf mice^38^. We note that the matching WT controls were not used to train the RRBS clock. Excitingly, both Snell Dwarf and GHRKO mice were predicted as significantly younger than the matching controls (**Figure 3f**). Thus, we conclude that both individual CpG and combined CpG TE clocks show an association with the health status of the individual and not only their chronological age.

**Figure 4:**
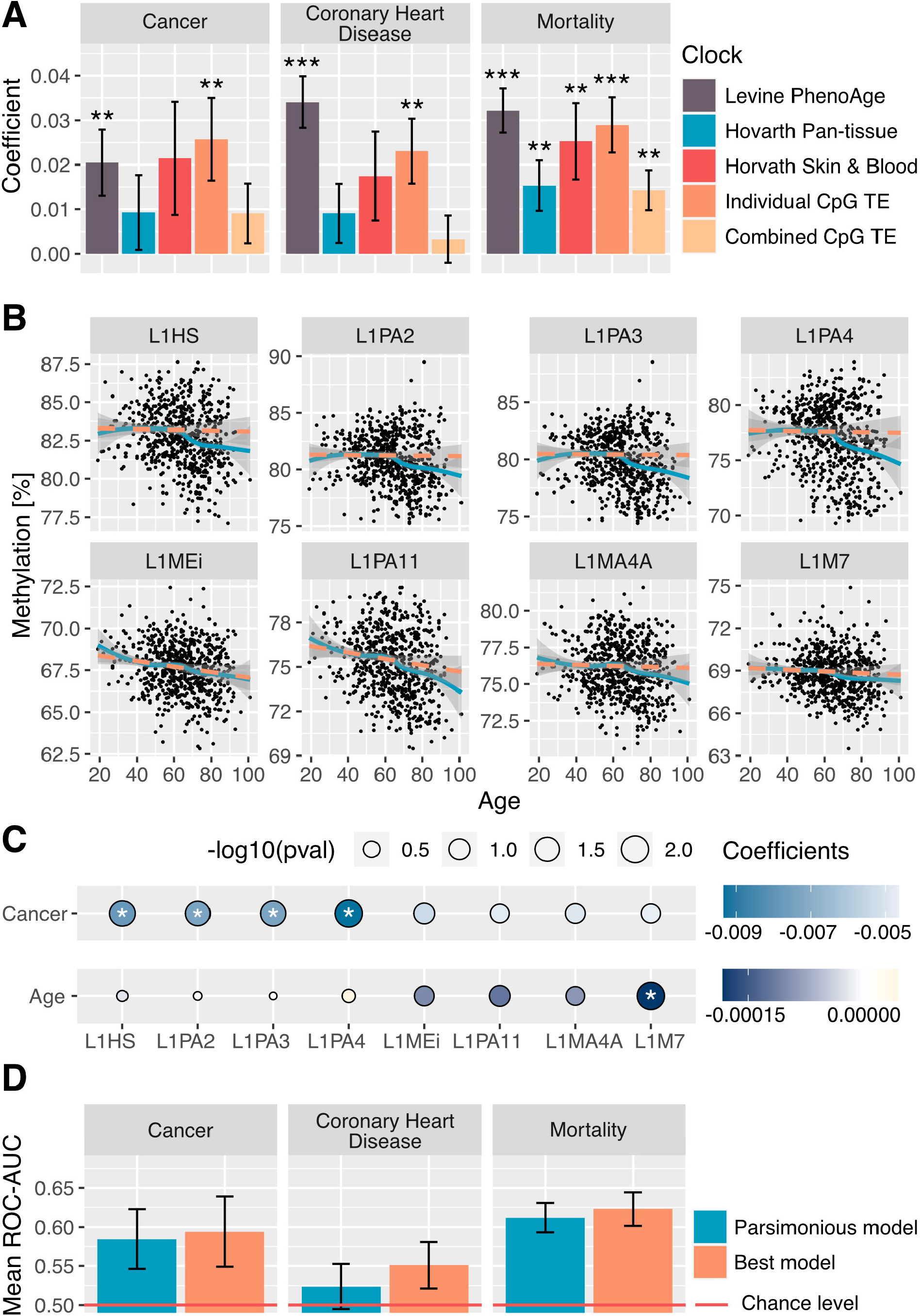
Association between TE clock acceleration, TE methylation loss and disease. (A) Association between age acceleration and risk of cancer, CHD and mortality according to the individual and combined CpG clocks in the WHI BA23 dataset. Results are benchmarked against state of the art chronological age clocks (Horvath Pan-tissue and Horvath Skin and Blood) and biological age clocks (Horvath PhenoAge). (B) Age trajectory of methylation at young L1s (first row) and old L1s with the largest coefficients in the combined CpG clock (send row). Data from GSE40279. Orange dashed line shows a linear fit, excluding patients over 65 years-old. Teal line shows a loess fit on the full age range. (C) Effect of cancer within 3 years and age on methylation of young and old L1s in the WHI data. (D) Performance of predictors of risk of cancer, CHD and mortality within 3 years. Best and parsimonious models are shown.

### Properties of young and old L1s as biomarkers

Finally, we investigated the TE families selected by our combined CpG clocks. Among the notable TE families we identified, only L1HS (genic) was chosen as a feature by human combined CpG clock, with methylation loss associating with increased age. However, several older L1 families were chosen with stronger coefficients (L1MEi, L1PA11, L1MA4A, L1M7 …). We found this puzzling, as we expected that the strong age association of younger L1s (L1HS, L1PA2, L1PA3 and L1PA4) would make them useful for age prediction. Thus, we investigated the exact trajectory of young L1 de-repression in greater detail (**Figure 4b**). We were surprised to see that young L1s had negligible methylation loss under the age of 65 and then rapidly lost methylation in older patients with a non-linear trajectory. In comparison, the older L1 families selected by our combined CpG predictor showed a more linear trajectory, and began demethylating at younger ages. This led us to suspect that older, “passively demethylating” TE families may be better predictors of chronological age, whereas methylation loss at younger TEs, in particular those with pathogenic potential, may be better predictors of disease risk. Thus, we modelled average methylation at young L1s (L1HS, L1PA2, L1PA3, L1PA4) and old L1s with large clock coefficients (L1MEi, L1PA11, L1MA4A, L1M7) as a function of age, this time including whether individuals would be diagnosed with any cancer within 3 years of sample collection (Methylation ∼ Age + AnyCancerIn3y, **Figure 4c**). We found that cancer was significantly associated with decreased methylation of young L1s, but not at older ones, although a trend was still present. Conversely, when accounting for cancer, age was associated with decreased methylation at older L1s but not at young ones. With this knowledge we trained predictors of cancer, CHD and mortality within the next 3 years solely based on young L1 CpGs (n=621) in the WHI data. These events were quite rare (cancer: n = 52, chd: n = 140, death: n = 39, total: n = 2175) making training challenging. Nonetheless, the resulting models had mild predictive ability (**Figure 4d**). Interestingly, while the mortality and CHD predictors were rather complex, even when choosing the optimal model with parsimony (Best mortality predictor: ncpg = 93, parsimonious mortality predictor: ncpg = 60; best CHD predictor: ncpg = 180, parsimonious CHD predictor: ncpg = 106, **data file 3**) the cancer predictors were remarkably simple, using only a handful of CpGs. The simplest model based predictions on just 2 CpGs: cg07575166, found in an intergenic L1HS 5’UTR, and cg26106149, located in a full length L1PA3 in an intron of FBXL4, a gene with no known role in cancer initiation. The more complex model used 5 more CpGs but assigned the most weight to the aforementioned 2.

## Discussion

In summary, we studied the age dynamics of TE methylation, finding that most TEs, from evolutionarily young, to ancestral ones, were likely to lose methylation during the course of aging. However, this tendency was accentuated for young L1 elements: L1HS, L1PA2, L1PA3 and L1PA4, and two LTR families: THE1A and THE1C. Local CpG density and youthful methylation have been previously reported to affect methylation drift rate during aging. The rate of methylation loss at most TEs was well described by those two factors, but this was not the case for young L1s. Thus, we hypothesize that most TEs have lost their regulatory sequences, and thus lose methylation passively. Conversely, young L1s are likely to still contain regulatory sequences that enable recruitment of activating epigenetic machinery. We next explored the use of TE methylation loss as biomarkers of age and disease. An age predictor based on individual CpGs found in TEs had remarkable accuracy, and showed associations with cancer and mortality comparable to PhenoAge. We generated additional predictors based on average methylation of TEs genome-wide, for both human blood methylation array data and multi-tissue mouse RRBS data. While less accurate than their individual CpG counterparts, these predictors were still satisfactory (r > 0.9) and showed associations with health status. We were surprised to see that these predictors did not mainly rely on young L1s despite their strong age association, prompting us to investigate the exact timing of young L1 derepression. We found that young L1s rapidly derepressed only after age of 65 and were otherwise very stable beforehand. This age coincides with the age of onset of many age-related diseases. Thus, we explored associations between loss of methylation and disease, finding that methylation loss at young L1s was associated with cancer but not age, while the opposite was true for the older L1s selected by the clock. Finally, we trained predictors for cancer, CHD, and mortality within 3 years of the methylation measurement, solely based on young L1 CpGs. The mortality and cancer predictors were mildly successful and, in particular, the cancer predictor made use of only 2 CpGs in young L1s. Future studies may investigate the mechanism behind this seemingly direct relationship. An obvious question is whether young L1 derepression is the cause or consequence of cancer. Indeed, both mechanisms are possible, as mutations of epigenetic machinery are common in cancer^39^. However, as the loss of CpGs was detected in the blood and was predictive of cancer events in other organs, it is possible that TE derepression may promote cancer by accelerating inflammation or by promoting other pathological processes through other non-cell autonomous mechanisms. Finally, loss of methylation at young L1s could be neither the cause nor the consequence of cancer, and instead both events could have common drivers. The clonal haematopoiesis is a likely suspect, as the most common mutation in clonal hematopoiesis is DNMT3A, a de-novo methyltransferase^40–42^, which may also contribute to the loss of methylation on TEs.

## Methods

### Datasets

We used 4 public human blood array datasets (GSE64495^43^, GSE40279^21^, GSE157131^44^, GSE147221^45^) to determine associations between age and TE methylation loss, and to train and validate the human age predictors. GSE87648^46^ was only included in predictor training and validation because it appeared to have an internal batch effect (determined by PCA). The WHI human blood dataset BA23 (https://www.whi.org/study/BA23) and related metadata were used to investigate relationships between TE clock age acceleration and risk of disease and mortality, and later investigate associations between young L1 methylation loss and disease. Mouse multi-tissue datasets GSE60012^47^, GSE93957^48^, GSE80672^38^ we used to train and validate the mouse age predictor. All data was used as pre-processed by the original authors with the exception of GSE60012, as the needed processed files were unavailable.

### Annotation of CpGs and repetitive elements

The coordinates of infinium array CpGs were obtained from the Illumina manifest. We used RepeatMasker to annotate repeats in GRCh37 and GRCm38 genomes. ChipSeeker^49^ was used to annotate the genomic context of CpGs.

### Statistics

Associations between age and Infinium array CpG methylation were determined using limma^29^, with the design ∼ age + sex + study. The fitted coefficients were used as methylation drift rates, whereas methylation at 20 years of age was calculated as intercept + coef * 20. Our fitting of expected age drift as function of CpG density and youthful methylation level employed a general additive model (gam) with covariates for CpG density within 100 bp of the CpG in question, the methylation of that CpG at 20 years of age, and the interaction of the two covariates (age_coef ∼ s(methylationAt20yo, bs = “cs”) + s(CpG_density, bs = “cs”) + s(methylationAt20yo, bs = “cs”, by=CpG_density)). Associations between age acceleration and mortality/disease risk were tested using a Cox regression model (coxph in R) with formula Surv(time-to-event, status) ∼ acceleration + age.

### Predictor training and validation

All predictors in this study are a form of elastic net, implemented by the glmnet R package. Age predictors use the gaussian family argument whereas the disease/mortality predictors use the binomial (logistic) family argument. Age predictions were evaluated by root mean squared error(RMSE), median absolute error(MAE) and pearson’s r. Disease/mortality predictions were evaluated by receiver operating characteristic area-under-the-curve (ROC AUC). Prior to training/predicting, we transformed ages using the same age transformation used by Horvath in the Pan-tissue^22^ and Skin & Blood^23^ clocks. Briefly, ages below the age of maturity (20 years for humans, 6 weeks for mice) were log transformed, to linearize the relationship between age and methylation in developmental stages. When sufficient samples were available, we validated our predictors by leaving out a portion of all data and an entire dataset (GSE64495) for testing, and training/choosing hyperparameters on the remainder of the data by cross-validation. When the number of samples was limited, we used nested cross-validation. Hyperparameters explored by grid search and selected to give the lowest cross-validation MSE (mean squared error) or ROC AUC, with the exception of the models we called “parsimonious” for which hyperparameters were selected to give the simplest model within 1 standard deviation of the best performance. Any individual with known health conditions or treatments were excluded from model training. The matching wild-type controls of GHRKO and Snell dwarf strains were also excluded from clock training, to have a fair comparison.

### Predictor benchmarking

We downloaded clock coefficients published with the original manuscripts. Ages were transformed (and inverse transformed) for prediction if required (Horvath Pan-tissue and Skin & Blood). All clocks were then applied to the same samples of GSE64495 and the WHI BA23 dataset. Clock features with missing values in the WHI BA23 (1.5% of all values) were imputed using the makeX R function.

### RRBS data processing

Raw reads were downloaded from SRA and trimmed using TrimGalore!^50^ with the --rrbs option. We aligned trimmed reads to the GRCm38 genome build using Bismark^51^ and quantified methylation with bismark_methylation_extractor and bismark2bedGraph.

## Acknowledgements

This research was supported by grants from the US National Institutes of Health to AS and VG, and Milky Way Research Foundation to VG. A.H.S. was supported by grant RF1AG074345 from the National Institute on Aging. The WHI program is supported by contracts from the National Heart, Lung and Blood Institute, NIH. The authors thank the WHI investigators and staff for their dedication, and the study participants for making the program possible. A listing of WHI investigators can be found at https://www.whi.org/doc/WHI-Investigator-Long-List.pdf.

## Figure legends

**Figure S1:**
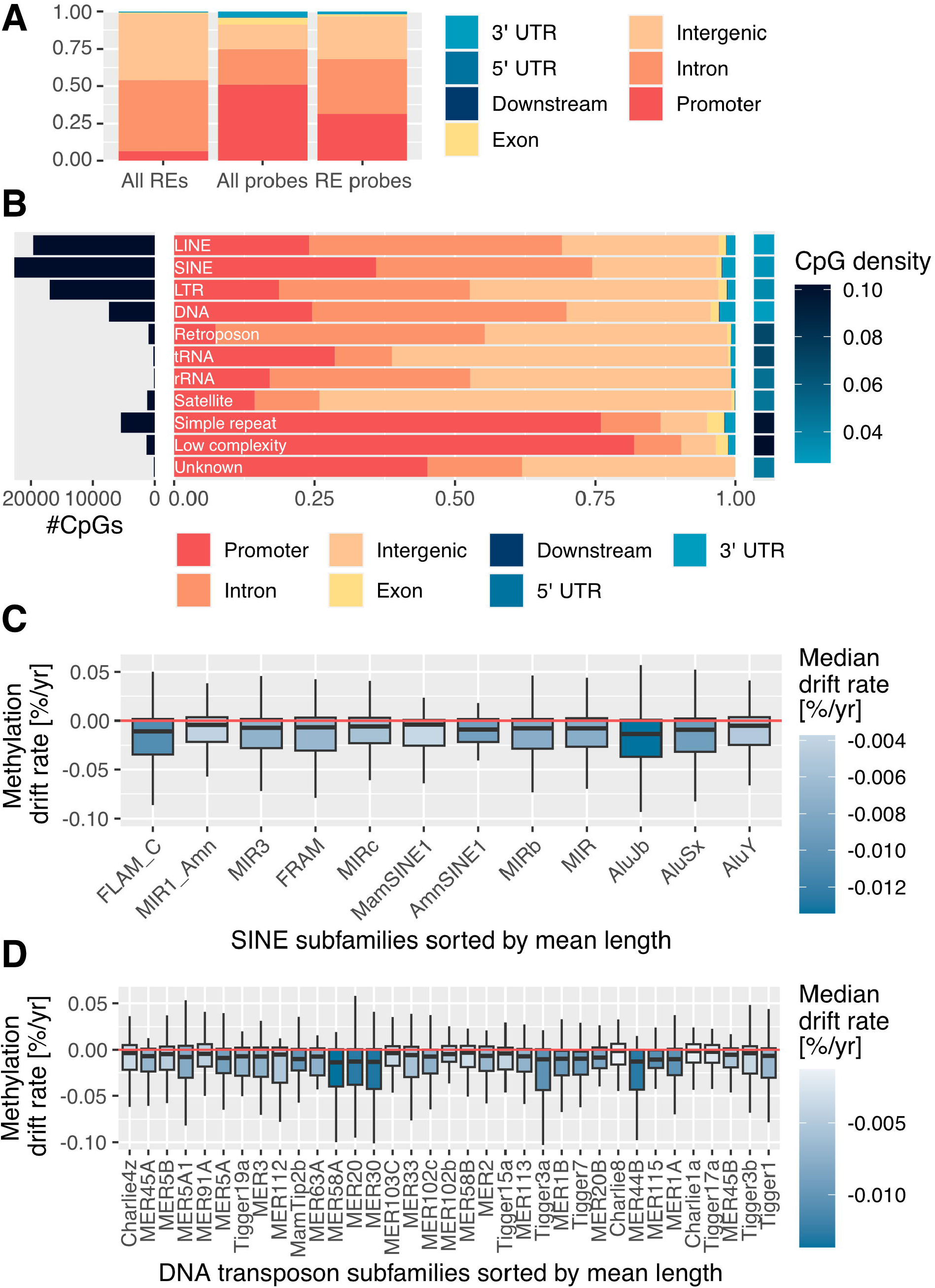
Genomic context of RE CpGs. Age trends of SINE and DNA transposon methylation. (A) Genomic context of all REs, all probes in the infinium array, RE probes in the infinium array. (B) Genomic context of infinium probes by major RE class. (C) Methylation drift rate of CpG in SINEs, grouped by family and sorted by average sequence length. Only families represented by 40 or more CpGs in the infinium array were shown. (D) Methylation drift rate of CpG in DNA transposons, grouped by family and sorted by average sequence length. Only families represented by 40 or more CpGs in the infinium array were shown.

**Figure S2:**
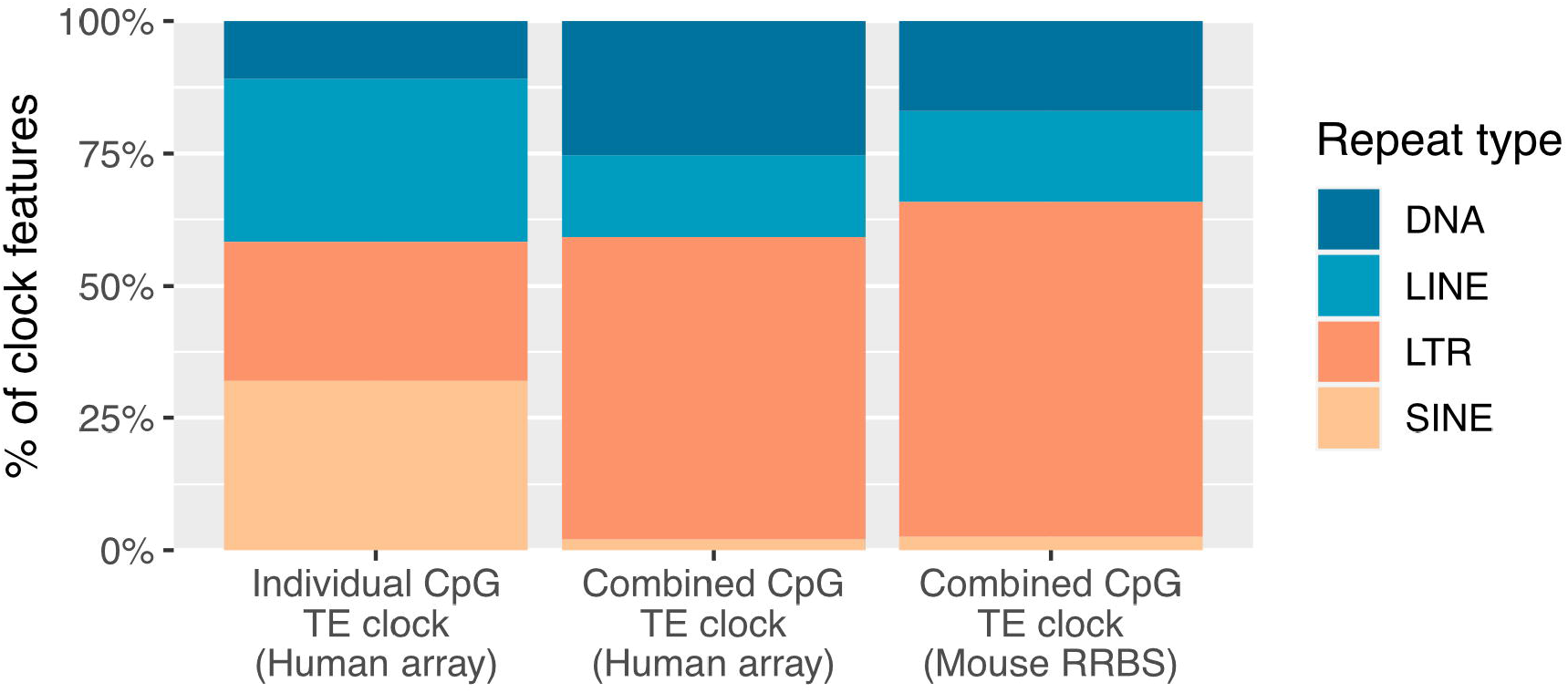
Composition of TE clocks by class.

